# Meta-Analysis of Transcriptomic Variation in T cell Populations Reveals Novel Signatures of Gene Expression and Splicing

**DOI:** 10.1101/727362

**Authors:** Caleb M. Radens, Davia Blake, Paul Jewell, Yoseph Barash, Kristen W. Lynch

## Abstract

Distinct T cell subtypes are typically defined by the expression of distinct gene repertoires. However, there is variability between studies regarding the markers used to define each T cell subtype. Moreover, previous analysis of gene expression in T cell subsets has largely focused on gene expression rather than alternative splicing. Here we take a meta-analysis approach, comparing eleven independent RNA-Seq studies of human Th1, Th2, Th17 and/or Treg cells to identify transcriptomic features that correlate consistently with subtype. We find that known master-regulators are consistently enriched in the appropriate subtype, however, cytokines and other genes often used as markers are more variable. Importantly, we also identify previously unknown transcriptomic markers that consistently differentiate between subsets, including a few Treg-specific splicing patterns. Together this work highlights the heterogeneity in gene expression between isolates of the same subtype, but also suggests additional markers that can be used to define functional groupings.

## Introduction

The adaptive immune response relies on the ability of T cells to detect foreign antigen and respond by carrying out appropriate functions, such as the secretion of cytotoxins or cytokines. Importantly, T cells are not a uniform cell type, rather it is now recognized that multiple subtypes of T cells are generated during development and/or an immune response based on the nature of the foreign antigen and/or the context in which the antigen engages with T cells (DuPage and Bluestone, 2016). In particular, subtypes of CD4+ T cells differ in the antigens they engage and in the nature of their response to antigen, resulting in the optimal functional response to various types of immune challenge. However, while the functional impact of CD4+ T cell subtypes is clear, the molecular characteristics that distinguish different subtypes is yet to be fully established.

Two of the best characterized T cell subtypes are the T-helper 1 (Th1), T-helper 2 (Th2) subsets of CD4+ T cells, each of which express signature cytokines. Th1 cells secrete the cytokine interferon gamma (IFNγ) to promote an innate immune response against viruses or cancer (Tran et al., 2014), while parasite-fighting Th2 cells secrete the cytokines interleukin (IL)-4, IL-5, and IL-13 (Pulendran and Artis, 2012). Another commonly studied CD4+ subtype are T-helper 17 (Th17) cells which secrete IL-17 and IL-23 to fight fungal infections (Harrington et al., 2005) or cancer. Importantly, an inappropriate balance of T cell subtypes not only reduces effectiveness of fighting pathogens but can also cause disease. Overabundance or activity of Th1 cells contribute to colitis (Harbour et al., 2015), Th2 cells contribute to asthma and allergies (Venkayya et al., 2002), and Th17 cells contribute to multiple sclerosis and other autoimmune diseases (Vaknin-Dembinsky et al., 2006).

In addition to expression of the signature cytokines mentioned above, T cell subsets have historically been defined based on the expression of a lineage-defining master transcription factor (Shih et al., 2014). The Th-specific master regulatory transcription factors include: T-box transcription factor TBX21 (T-bet) for Th1 (Szabo et al., 2002), GATA-binding protein 3 (GATA3) for Th2 (Zhang et al., 1997; Zheng and Flavell, 1997), and RAR-related orphan receptor gamma (RORγ) for Th17 (Ivanov et al., 2006). However, using these two key factors, a master regulator and signature cytokines, to define T cell subsets is increasingly appreciated to be too simple to adequately explain the breadth and plasticity that has been observed for T cell populations (DuPage and Bluestone, 2016). For example, another common T cell subtype are regulatory T cells (Treg), which suppress immune responses. Treg suppressive function was found to depend on the master regulator Forkhead Box P3 (FOXP3) (Zheng and Rudensky, 2007), but there are no well-defined signature cytokines for Treg cells. Treg cells produce the *TGFB*, IL-10, or IL-35, which are critical anti-inflammatory cytokines, but not all of these cytokines are produced by all Treg cells (Shih et al., 2014). Moreover, core signature cytokines and master regulators such as IFNγ and T-bet have been observed in Th17, Th22 and other hematopoietic cells (Shih et al., 2014).

Another traditionally oversimplified characteristic of Th cells is their ability to sense and migrate to specific chemokines by expressing distinct and specific chemokine receptors under the direction of T cell subset master regulators (DuPage and Bluestone, 2016). Thus, T cell subsets are often identified *ex vivo* by extracellular expression of chemokine receptors including: *CXCR3* for Th1, *CCR4* for Th2, *CCR6* for Th17 and *IL2RA* for Tregs. However, T cells have shown remarkable plasticity in their ability to polarize between distinct T cell subtypes (Bending et al., 2009; DuPage and Bluestone, 2016) and transitionary T cells exist that simultaneously express chemokine receptors from two subsets (Cohen et al., 2011). Therefore, it is clear that better descriptions are needed of the molecular differences that define functionally distinct T cell subpopulations.

An additional complication to identifying molecular markers that differentiate one CD4+ subtype to another is that methods to isolate populations vary widely across the field. One approach to purifying T cell subsets is the use of antibodies to chemokine receptors or other extracellular marker proteins to isolate specific T cell subsets using flow cytometry or magnetic beads. However, these methods are limited by the above-mentioned variability and overlap in expression of these proteins. By contrast, an alternate method to enrich for T cell subsets is polarizing naïve CD4+ T cells toward distinct phenotypes *in vitro* with various cytokine cocktails. While these cytokine cocktails are meant to mimic the environment that promotes the development of each subtype, the exact conditions used vary from one laboratory to another, thereby also inducing variability between studies.

Here we take a meta-analysis approach across a range of studies that have isolated and defined T cell subsets by a variety of methods. Using this approach, we sought to determine if there is a set of transcriptomic features that are consistent for particular subset independent of the experimental methodology. In addition, while most previous studies have focused on differences in gene expression between T cell populations, we also investigated the extent to which alternative splicing contributes to transcriptomic variation between subsets. Recent studies have revealed widespread and co-regulated alternative splicing early in CD4+ T cell activation in the absence of polarizing cytokines (Ip et al., 2007; Martinez et al., 2015a; Martinez and Lynch, 2013). Alternative splicing is a ubiquitous mRNA processing step whereby, for example, genes express isoforms including or lacking an alternative exon. Such splicing variations often results in different protein isoforms which can have disparate functions (Braunschweig et al., 2013). While a few studies have reported differential splicing or expression of splicing regulatory proteins in particular T cell subsets in mice (Middleton et al., 2017; Stubbington et al., 2015), a comprehensive comparison across human T cell subsets has not been reported. By comparing RNA-Seq data across eleven independent studies we identify a set of ∼20-50 genes whose expression is well-correlated with distinct T cell subsets. Notably, these include some, but not all, of the genes encoding the cytokines and receptors typically used to define T cell subsets, as well as some additional genes that have not previously been considers indicators of cell fate. We also find a smaller set of genes for which splicing patterns correlate with cell subtype; however, splicing seems to be less definitively regulated in a subtype-specific manner than transcription. Together, these data are consistent with recent models arguing for more elaborate and nuanced definitions of T cell populations. Moreover, our data provide critical information as to the underlying biologic differences between CD4+ subsets and what combination of molecular markers may be best used to define these populations.

## Results

### Selection of datasets and RNA-Seq analysis pipeline

Given the extensive variability in the methods used in the field to generate and define T cell subsets we were interested in determining how much molecular variation exist in the resulting cell populations and whether there are transcriptomic signatures that robustly identify T cell subsets regardless of experimental conditions. To carry out a meta-analysis of transcriptomic variation in T cell subtypes, we identified RNA-Seq experiments in the NCBI-GEO and EMBL-EBI-ArrayExpress databases that were from polyA-selected RNA samples derived from CD4+ T cells purified from the blood of healthy humans and had at least two out of three of the following types of samples: naïve (no stimulation), Th0 (stimulated in the absence of polarizing cytokines), or specific T cell subsets. We also confirmed the quality of the datasets by requiring that all samples had greater than 60% uniquely mapped reads and no non-human overrepresented sequences. Lastly, we performed principle component analysis of samples by gene expression to confirm that in each study samples clustered by cell type (Supplemental Figure S1). In total, 10 publicly available datasets, plus one in-house dataset (Bl, GSE135118), met these inclusion and quality control criteria (Figure 1A and Supplemental Table S1). These 11 datasets include multiple replicate samples of naïve and Th0 CD4+ cells, as well as Th1, Th2, Th17 and Treg subpopulations (Figures 1A and 1B). Importantly, these datasets encompassed samples of specific T cell subsets obtained using one of two general approaches: (1) sorting cells from whole blood using known extracellular protein markers, or (2) *in vitro* polarization of naïve CD4+ T cells towards Th1, Th2 or Th17 cell fates with specific cytokine cocktails. Datasets that generated T cell populations by *in vitro* polarization obtained naïve precursors from either adult whole blood or neonatal cord blood and used distinct cytokine combinations (Figure 1A). A detailed description of the experimental conditions and RNA-Seq technical specifications for each dataset is shown in Supplementary Table S1.

**Figure 1:**
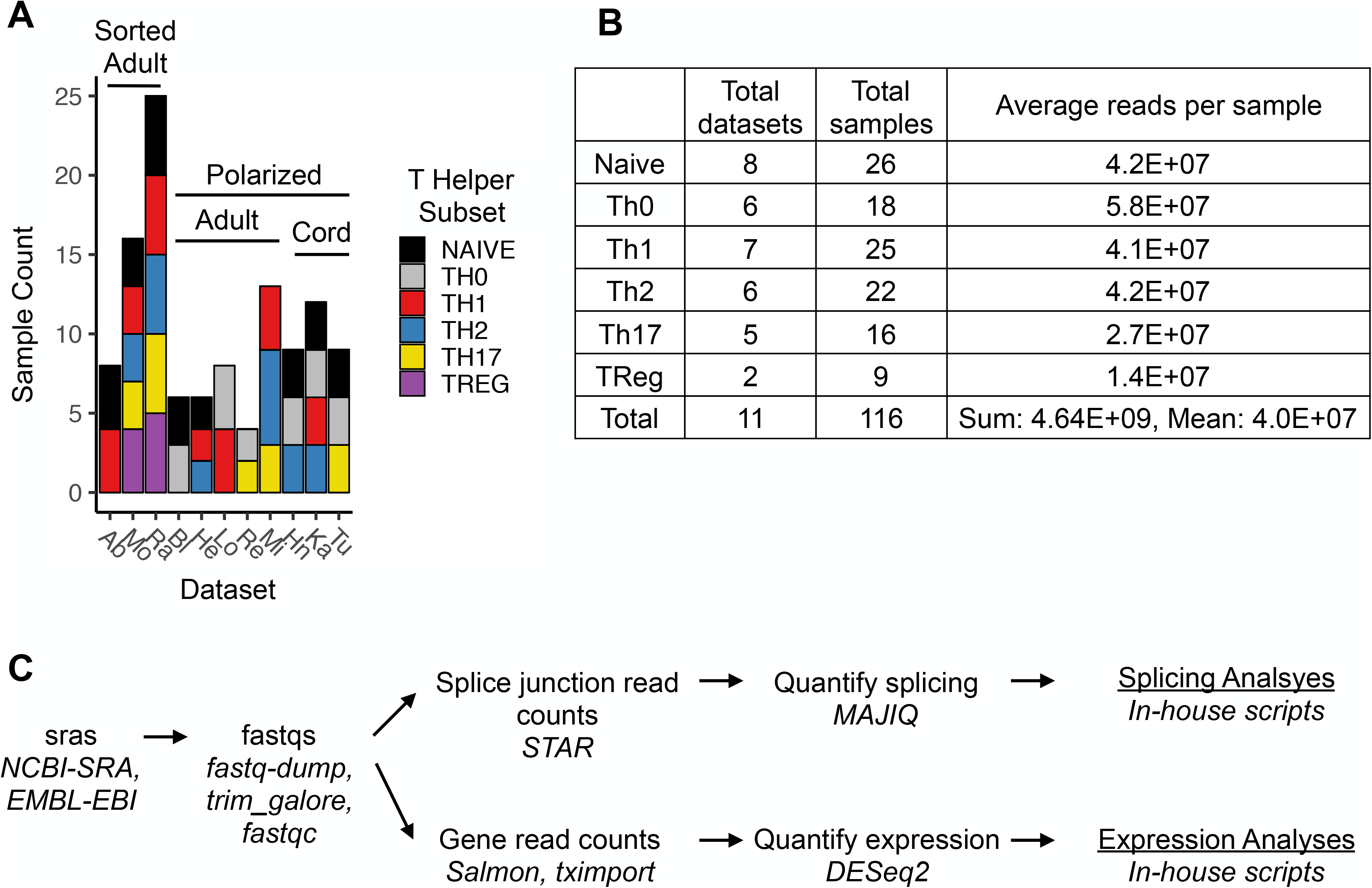
Datasets and analysis pipeline used for meta-analysis. (A) Datasets used in this study and the breakdown of samples and isolation methods for each dataset. Sorted: samples sorted from blood, Polarized: samples *in vitro* polarized, Adult or Cord: cells derived from adult or cord blood. Ab (Abadier et al., 2017), Mo (Monaco et al., 2019), Ra (Ranzani et al., 2015), Bl (this study), He (Hertweck et al., 2016), Lo (Locci et al., 2016), Re (Revu et al., 2018), Mi (Micossé et al., 2019), Hn (Henriksson et al., 2019), Ka (Kanduri et al., 2015), Tu (Tuomela et al., 2016). For further detail see Supplemental Table S1. (B) Total number and quality of datasets for each T cell subtype used in this study. (C) Pipelines used to process RNA-Seq data and quantify gene expression and splicing variation.

To begin to look for common patterns of gene and isoform expression in the above datasets, raw RNA-Seq data from all samples were uniformly processed by trimming low quality base calls and adaptor sequences and aligning reads to the hg38 genome (Figure 1C). Gene expression was then quantified with the Salmon and DESeq2 algorithms, while local splicing variations were quantified with the MAJIQ algorithm (Figure 1C). Importantly, all but two of these datasets used a donor-paired experimental design (i.e. multiple cell types were isolated or derived from each given donor), allowing us to directly compare transcriptome profiles within T cell subsets from an individual donor. RNA-Seq-based genotyping was used to confirm all sample pairings, as well as to identify sample pairings in those studies in which such information was not given (see STAR methods).

### Master regulators are consistently expressed in a subtype-specific manner, while other common markers are not

In order to determine the validity and robustness of both the datasets and our analysis pipeline, we first assessed the expression of well-accepted signature genes of Th1, Th2, Th17, and Treg cell populations across the various subpopulations. Although the absolute expression of previously-described “master regulators” for Th1 (*TBX21*), Th2 (*GATA3*), Th17 (*RORC*) and Treg (*FOXP3*) varied from study to study, within any given study the majority of the Th1, Th2, Th17, and Treg samples expressed their respectively known master regulators more highly than any other cell type (Figure 2A). For example, 25 of the 25 Th1 populations across all seven Th1-containing datasets, expressed the Th1 master regulator *TBX21*, as or more highly than other cells in the same study (Figure 2A, top panel). Similarly, 21 of 22 Th2 samples highly expressed the Th2 master regulator *GATA3* (Figure 2A, second panel), and all of the Th17 and Treg samples were enriched for their respective master regulators *RORC* and *FOXP3* (Figure 2A, bottom two panels). As emphasized above, each study analyzed here used different protocols to obtain T Helper cell RNA-Seq samples. Therefore, these enrichment data demonstrate that master regulator gene expression clearly segregates Th1, Th2, Th17 and Treg populations independent of experimental method.

**Figure 2:**
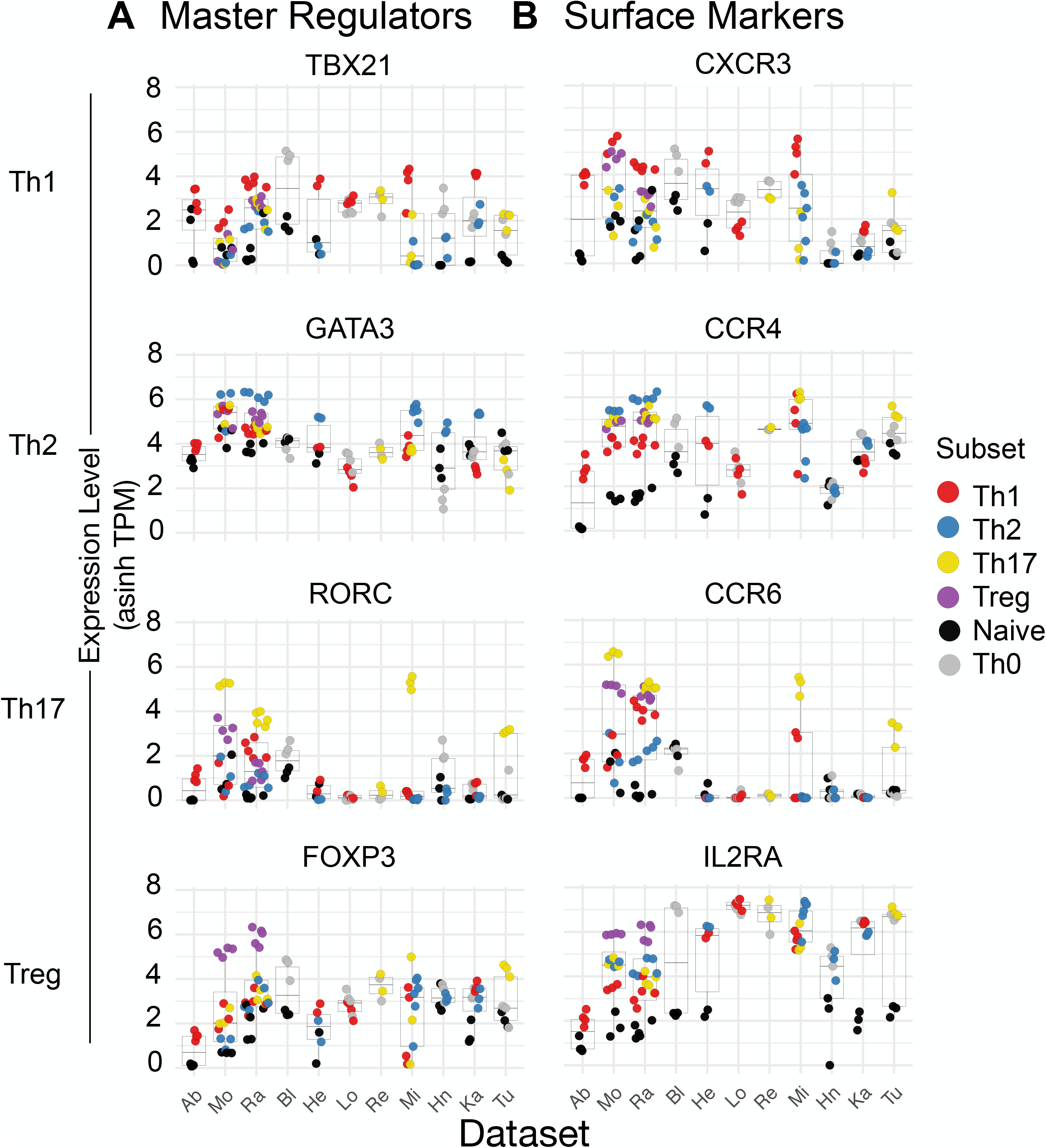
Expression of classical master regulators and cell surface markers across T cell subsets. Expression of signature genes typically used to delineate T Helper cells, including genes encoding (A) master transcription regulators and (B) cell-surface proteins, across all datasets. Each subplot represents a distinct gene, while each column in a specific subplot is a distinct study. Studies are listed at bottom and ordered as in Figure 1A. Cell type listed on left in bold is the subtype expected to be enriched for the two genes plotted on that row. For each study the mean and distribution of the given gene is displayed as a box plot, with the values for each individual sample represented as a dot. Dots are colored by cell subtype. Note that not all studies contain all cell types, as described in Figure 1A. Expression values shown as the inverse hyperbolic sine (asinh) of transcripts per million (TPM).

Interestingly, unlike the clear expression patterns of master regulators, several extracellular markers commonly used to isolate T Helper cell subpopulations showed less clear Th1, Th2, Th17, and Treg-specific expression patterns (Figure 2B). The Th1 extracellular marker *CXCR3* was preferentially expressed six out of seven datasets; however, in one dataset (Lo), Th1 cells express less *CXCR3* than Th0 cells. This variability in expression cannot be explained by method of cell isolation as Lo used similar methods as other studies that do show Th1-specific expression of *CXCR3* (*in vitro* polarized, He and Si). Similarly, *CCR4* only was enriched in Th2 cells in only three of the six datasets that include these cells (Mo, Ra, and He), while the Th17 extracellular marker *CCR6* showed Th17-specific expression patterns in only 9/16 samples across four out of five datasets. By contrast, the Treg extracellular marker *IL2RA* was highly expressed in 9/9 Treg samples from both datasets with Treg samples.

We also find significant variability in the extent to which T cell subtype samples expressed their expected signature cytokines (Figure 3). Almost all of the Th1 samples (22/25) do express the Th1-specific cytokine *IFNG* more highly than other cells types in the same study. By contrast, we observe no enrichment of the Treg-associated cytokines *TGFB* or *IL10* in Treg cells relative to others (Figure 3). Th17 and Th2 cells show some bias in expression of their associated cytokines, *IL4/5/13* and *IL17A/F* respectively, over other cell types, but only approximately half of the individual Th17 or Th2 cell samples express higher levels of these cytokines than other cells in the same studies (Figure 3). Taken together, the above analysis of genes previously associated with Th1, Th2, Th17, and Treg populations reveals significant variability of all but the “master regulator” genes. This raises concerns about the validity of the commonly-used cytokines and cell surface receptors as consistent and reliable identifiers of T cell subtypes. Moreover, the lack of correlation between the mRNA levels of “master regulators” and other expected cytokines and receptors suggests that the master regulators are not sufficient to induce the full program of gene expression that are often assumed to be associated with the identity and function of T cell subtypes.

**Figure 3:**
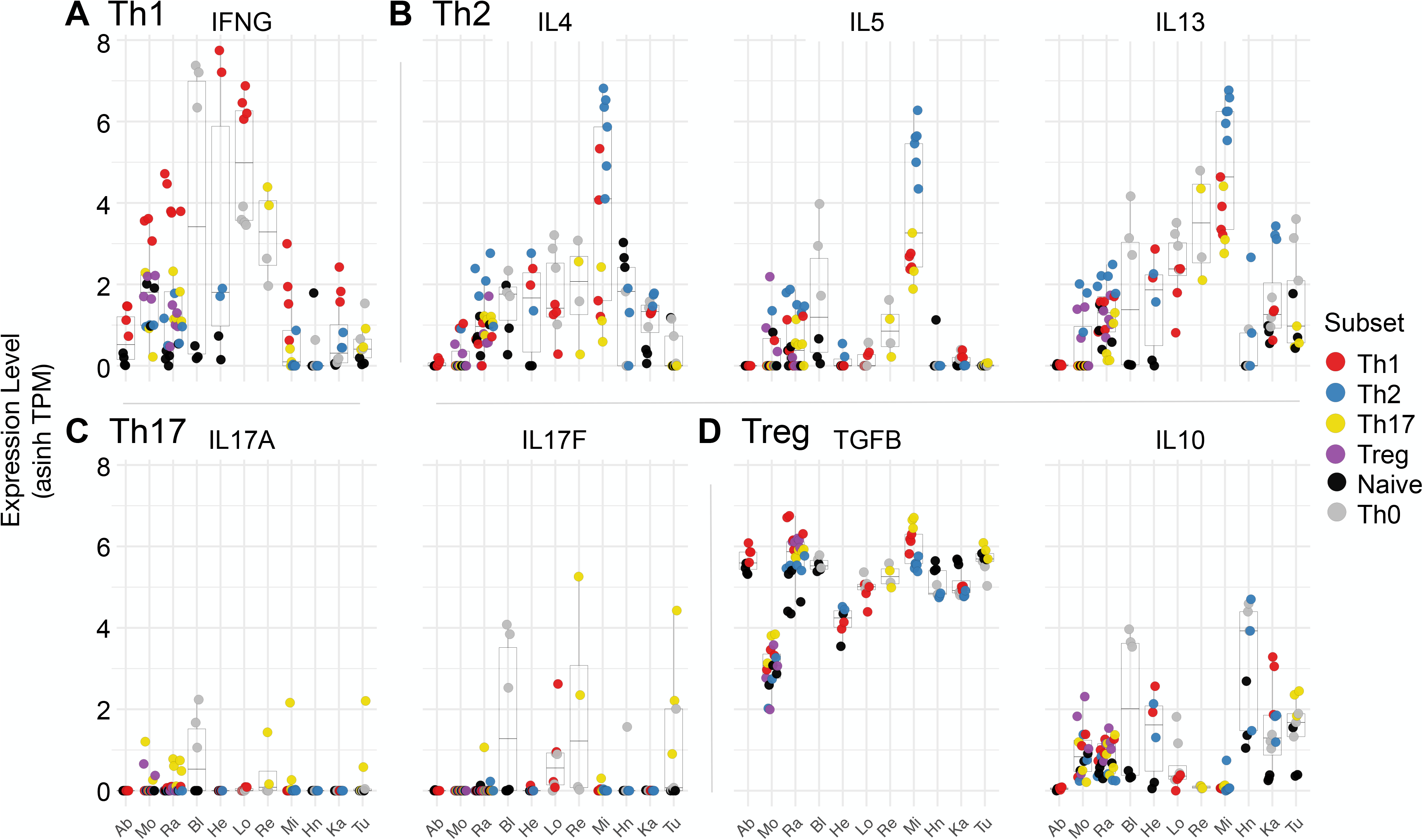
Expression of cytokines across T cell subsets. Expression of cytokines associated with (A) Th1, (B) Th2, (C) Th17 or (D) Treg cells, across all datasets. Each plot represents a distinct gene, while each column is a distinct study. Studies are listed at bottom and ordered as in Figure 1A. For each study the mean and distribution of the given gene is displayed as a box plot, with the values for each individual sample represented as a dot. Dots are colored by cell subtype. Note that not all studies contain all cell types, as described in Figure 1A. Expression values shown as the inverse hyperbolic sine (asinh) of transcripts per million (TPM).

### Th1, Th2, Th17, and Treg populations highly express core sets of genes

Given the relatively limited consistency of expression of genes expected to correlate with T cell subtypes, we next asked if other genes might be consistently enriched in Th1, Th2, Th17, and/or Treg cells regardless of variation in culture conditions and cellular sources. Specifically we mined the full gene expression data as determined by DESeq2 (Figure 1C) for genes “reliably expressed” in a given T cell subset, which we define as genes that are significantly more highly expressed in a given subset relative to others, in at least two of the studies analyzed (see Methods). 555 unique genes were reliably higher in at least one T cell subset over another across multiple datasets (Supplemental Table S2). These 555 subset-associated genes were then ranked by the number of datasets in which they were enriched, followed by the fold difference in expression between the subset analyzed versus others (Supplemental Figure S2).

Notably, by this ranking, the top 20 most associated genes for each T Helper subset (Figure 4) include the “master regulators” *TBX21* for Th1, *GATA3* for Th2, *RORC* for Th17, and *FOXP3* for Treg. In addition, consistent with the analysis of specific cytokines and chemokine receptors in Figures 2 and 3, the top Th1-associated genes include *CXCR3* and *IFNG*, the top 20 Th17-associated genes include *IL17A/F* and *CCR6*, and *IL2RA* is among the top 20 Treg-associated genes (Figure 4). By contrast, none of the Th2 signature cytokines or receptors genes (*IL-4/5/13* and *CCR4*) are significantly enriched in Th2 cells compared to the other populations surveyed (Figure 4).

**Figure 4:**
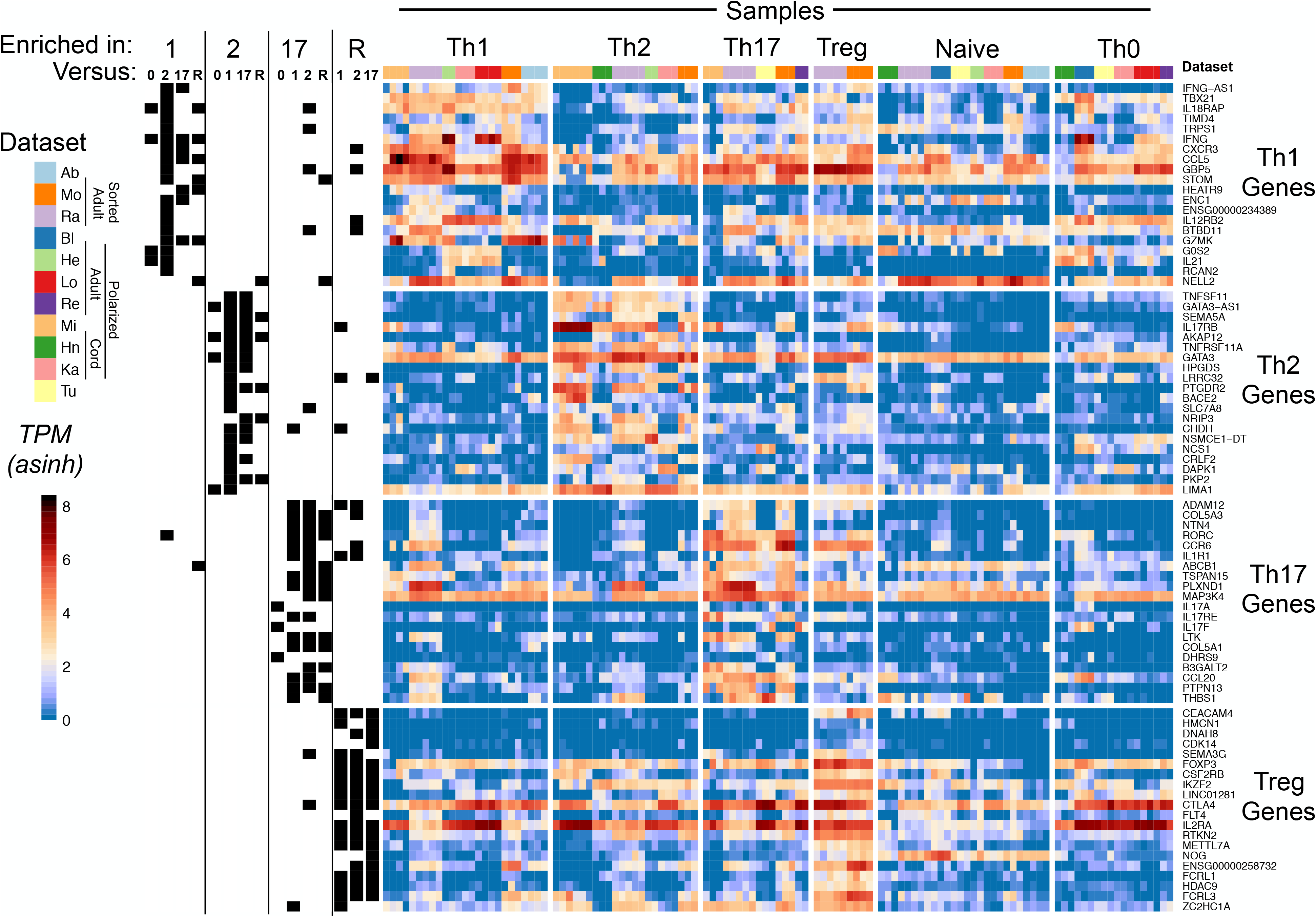
T cells consistently express core set of genes in a subset-specific manner. Reliably expressed T Helper genes (as defined in STAR Methods), ranked by number of supporting datasets and log2FC differences. Leftmost segment indicates whether a given core T Helper gene was reliably more highly expressed in a given subset over others. All differential expression analyses were performed between samples from the same dataset. The datasets with Treg samples lacked Th0 samples, so there is no “enriched in Treg vs Th0” column. Genes that exhibit enhanced expression between one subtype and multiple others are indicated by black boxes in multiple columns across a single row. The heatmap shows inverse hyperbolic sine (asinh)-transformed TPMs. Each column is a sample, and samples are grouped by cell type and author. Gene names are on the right.

Interestingly, our identification of subset-associated genes also highlights additional genes that may contribute to the function of particular T cell subsets and/or be useful as markers for subpopulations. Importantly, the expression of many of these genes is independent, at least at steady state, from the expression of the master regulator, as indicated by limited correlation in the expression levels of the genes in Figure 4 with the corresponding master regulators (Supplemental Figure S3). We emphasize that the genes highlighted in Figure 4 are consistently enriched in a T cell subset regardless of purification method or cell source, and thus are likely genes that are intimately tied to the biology of each subset. For example, *SEMA5A, IL17RB, AKAP12* and *HPGDS* have all previously been implicated in Th2 cell activity, and are both among the top 20 Th1-associated genes (Angkasekwinai et al., 2007; Lund et al., 2007; Mitson-Salazar et al., 2016; Zhang et al., 2013b); while *TNFSF11* (aka RANK Ligand), *TNFRSF11A* (aka RANK), and *LRRC32* are also among the highly Th2-associated genes, but it is unknown what role these genes play in Th2 biology. Notably *TNFSF11* and *SEMA5A* encode for cell-surface-exposed proteins and are more consistent Th2 cell markers at the mRNA level than *CCR4* or *IL4/5/13* (Figure 4), suggesting that TNFSF11 and SEMA5A proteins might have utility in isolation of Th2 cells by flow cytometry. Similarly, many of the genes enriched in Th1, Th17 and Treg cells have also been implicated in the biology of these cells, though not frequently used as markers for these subsets (see Discussion). Taken together we conclude that surveying a combination of differentially expressed genes may be more informative for defining and studying T cell subsets than simply relying on one or two markers.

### Analysis of local splicing variations reveals Treg-biased isoform expression

Given our success in identifying genes whose expression is highly associated with specific T cell subsets, we next asked the question of whether particular splicing patterns (i.e. mRNA isoforms) of certain genes was also correlated with T cell subtype. Strikingly, by contrast to the results with gene expression, we find very few instances in which a observe consistent differences in splicing patterns in a T cell subtype-specific manner (Supplemental Table S3 and Figure 5A). This lack of subset-specific splicing events is also contrary to the widespread changes in splicing we and others have previously observed between naïve and Th0 cells (Martinez et al., 2015b; Martinez and Lynch, 2013; Martinez et al., 2012). The few instances of subset-specific splicing that we can detect across datasets are cases of modest differential isoform expression in Treg cells versus Th2 or Th17 cells (Figure 5A). These splicing events represent all standard classes of splicing patterns (Figure 5B) and occur in genes that do not display any differences in overall expression (Supplemental Table S2 and Figure 5C). Therefore, the isoform differences that do exist between T cell subtypes are not readily detected in typical gene expression profiling.

**Figure 5:**
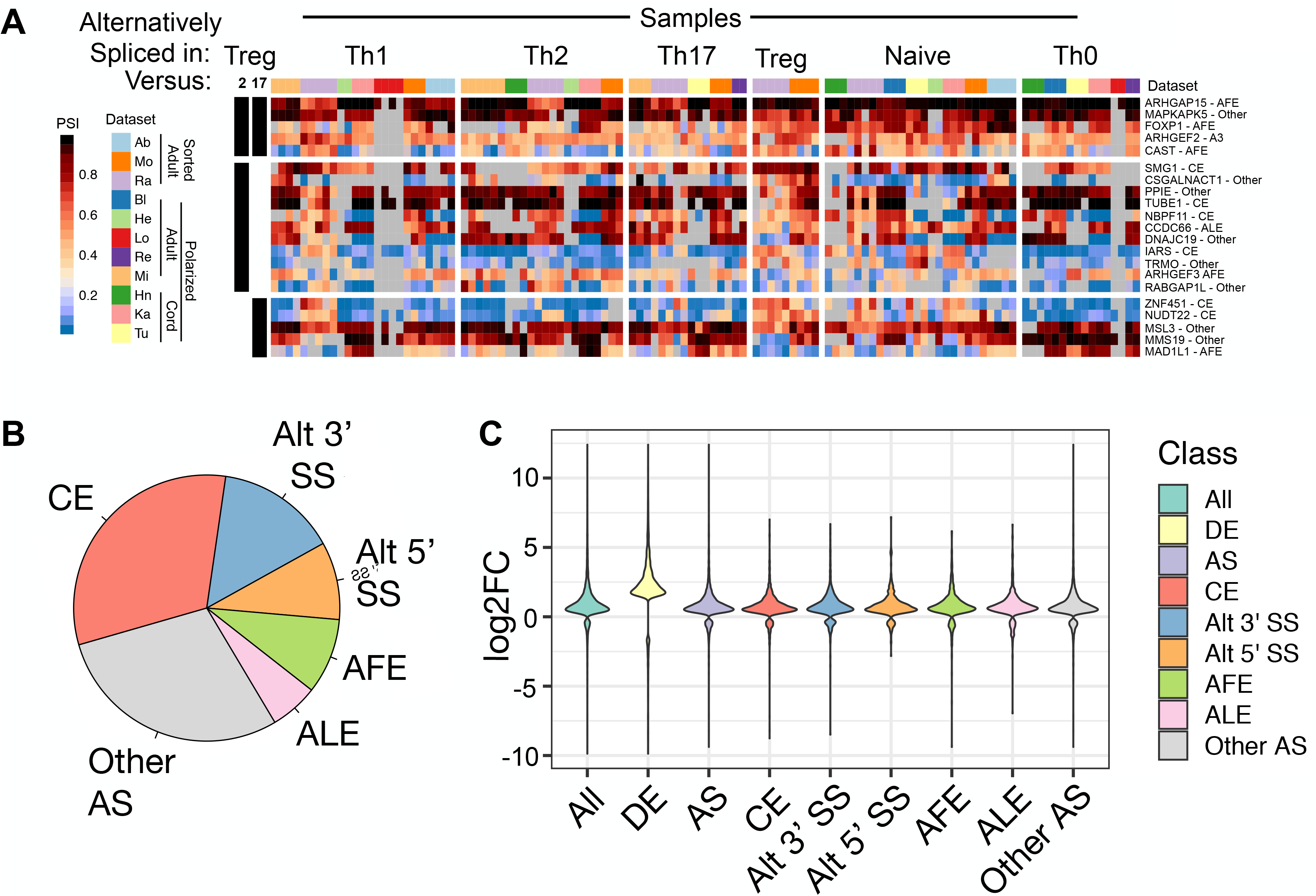
Only a limited number of splicing events are consistently regulated in a subset-specific manner in T cells. (A) Heatmap of PSI value for splicing events that show reproducible differences between Treg cells and Th2 and/or Th17 cells. Leftmost segment indicates comparisons that met significance threshold. Significant differences are based on comparison between samples from the same dataset (Ra and Mo have Treg), but PSI values are shown for all samples. Each column is a sample, and samples are grouped by cell type and author. Gene names and splicing event are on the right. (B) Pie chart of splicing events identified as differentially spliced between all T Helper subsets comparisons. Categories include cassette exons (CE), alternative first exons (AFE), alternative 3’ss (Alt3’ss), alternative last exon (ALE), or alternative 5’ss (Alt5’ss), or other non-defined patterns of splicing (Other AS). (C) Change in expression (Log2FC) in all genes that are differentially expressed (DE) or alternatively spliced (AS) between two cell subtypes studies as compared to all genes (ALL). The change in expression of each type of splicing event is also shown as for panel B.

Although the data in Figure 5 suggests that alternative splicing is not a general determinant of T cell identity, we do note that some of the observed Treg-biased splicing events are in genes linked to cytokine expression and immune function and thus are potentially of interest for future studies. For example, *ARHGEF2* exhibits differential use of alternative 3’ splice sites at the beginning of exon 7 in Treg cells versus Th2 and Th17 cells (Figure 5A and 6A). *ARHGEF2* encodes GEF-H1, a Rho guanine nucleotide exchange factor (Rho-GEF) that is involved in the response to intracellular pathogens and is required for expression of IL1β and IL6 (Wang et al., 2017). Tregs generally express more of the isoform that uses the distal 3’ splice site than Th2 or Th17 cells (Figure 5A, 6A). Use of this distal 3’ splice site results in the removal of a single alanine residue in the linker between the microtubule binding domain and the enzymatic Dbl-homology domain and has been shown to correlate with loss of RhoA-enhancing activity by GEF-H1 (Chen et al., 2019). A related Rho-GEF, *ARHGEF3*, is also somewhat differentially spliced between Treg and Th2 cells in that a proximal alternative first exon is favored in Tregs versus Th2 cells, thus altering the first 32-38 amino acids of the encoded protein. While functional differences have not been identified between the isoforms with distinct N-termini, ARHGEF3 has been linked to activation of RhoA in myeloid development (D’Amato et al., 2015).

**Figure 6:**
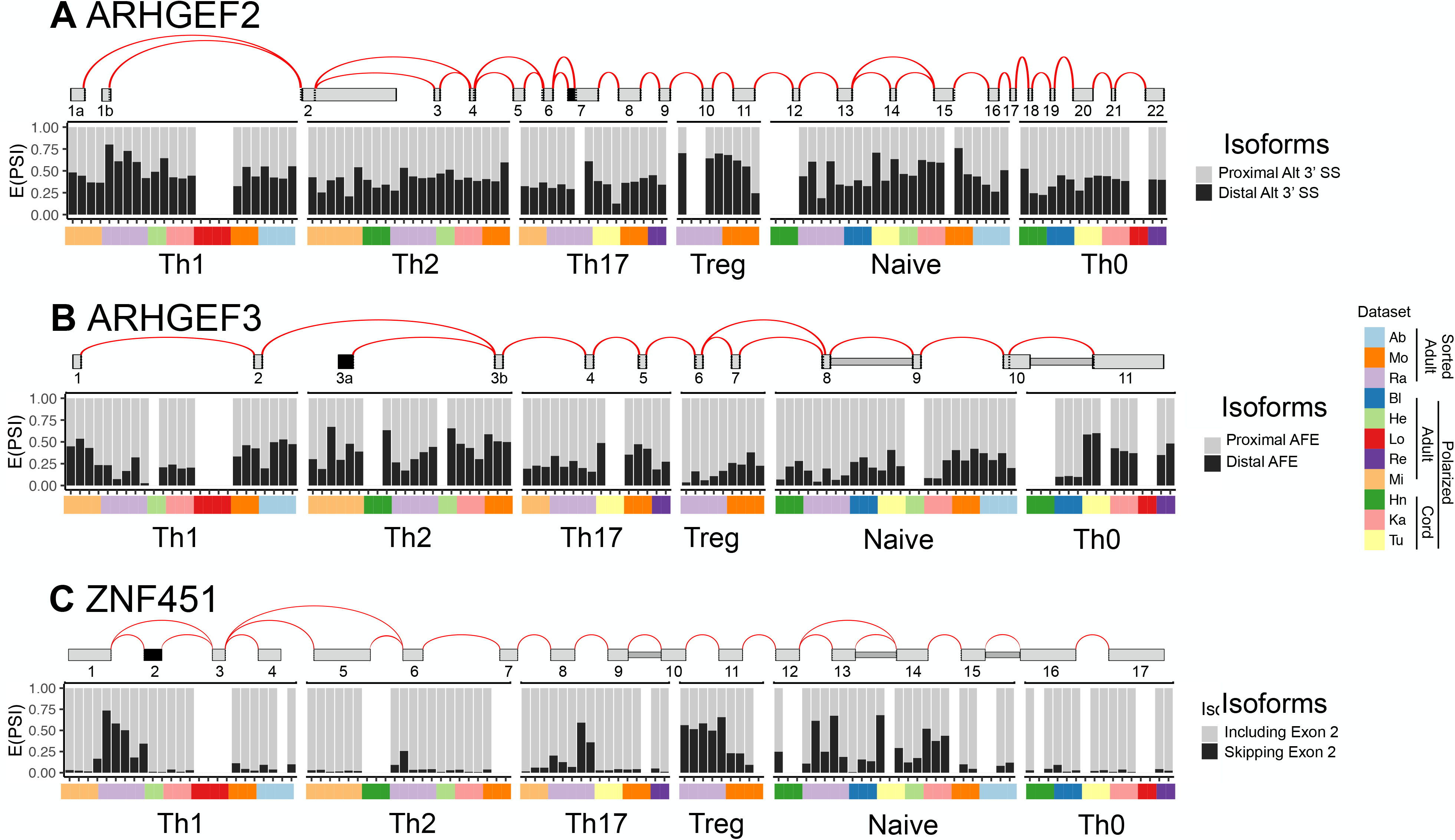
Treg-biased splicing events are predicted to alter protein function. Details of the splicing patterns for (A) ARHGEF2, (B) ARHGEF3 and (C) ZNF451 that exhibit some Treg bias (top) and the quantification of the variable events across samples (bottom).

Finally, a particularly interesting case is *ZNF451*, which exhibits preferentially skipping of an in-frame exon 2 in Treg cells as compared to Th17 cells (and perhaps also Th2 cells, although not scored as significant due to limited read count). ZNF451 is a SUMO E3 ligase and has also been shown to physically interact with Smad4 to repress its activation of TGF-β signaling (Cappadocia et al., 2015; Feng et al., 2014). Notably, exon 2, encodes part of the domain required to recruit SUMO to substrates (Cappadocia et al., 2015). While the role of the E3 ligase activity of ZNF451 to TGF-β signaling has not been defined, the differential expression of exon 2 in Treg cells suggests that this splicing difference may be a mechanism to differentially regulate TGF-β signaling in T cell subsets.

## Discussion

We and others have previously shown extensive changes in both gene expression and alternative splicing upon initial activation of naïve T cells (Martinez and Lynch, 2013). Specific changes in both gene expression and splicing have also been observed in discrete T cell subpopulations and assumed to be a consequence of polarizing conditions. However, since much variability exists in how T cell subsets are differentiated and defined, we sought to use a meta-analysis approach to investigate the variability of gene expression across different studies as well as to identify common transcriptomic signatures that are robust across all experimental definitions of specific T cell subsets. Importantly, while our analysis does confirm enriched expression of subtype-specific master regulators, we find that many other genes markers commonly associated with Th1, Th2, Th17 or Treg cells show limited predictive power to differentiate subtypes. On the other hand, our analysis also uncovered a handful of genes previously unknown to be regulated in a cell-type specific manner that show significant enrichment in one T cell subset compared to others. Finally, we show that while alternative splicing appears to play a limited role in shaping T cell differentiation into subsets, a few exceptions to this rule may exist especially in Treg cells.

Using a pair-wise comparison method we were able to identify ∼860 genes that exhibit differential expression between at least two T cell subsets (Figure 4, Supplemental Table S3). The top ten Th1 reliably expressed genes were mostly previously known to be important for Th1 biology, including the master regulator *TBX21* (Szabo et al., 2002), the Th1-associated cytokines *IFNG* and *CXCR3* (DuPage and Bluestone, 2016), the IFNG enhancer *IFNG-AS1* (Collier et al., 2014), as well as *IL18RAP* (Jenner et al., 2009), *TIMD4* (Nakajima et al., 2005) and *GBP5* (Lund et al., 2007). Other Th1-reliable genes included *TRPS1* and *CCL5*, which are not known to be important for Th1 cell populations specifically but are implicated in Th17 (Yosef et al., 2013) and memory cells (Swanson et al., 2002), respectively. *STOM* was also reliably expressed in Th1 samples, but it is not known what role, if any, *STOM* plays in T cell biology. *CCL5* and *TIMD4* encode for cell-surface-exposed proteins, so it would be especially interesting if these proteins could be used for isolating or quantifying Th1 cell populations.

Similarly, the top ten reliably expressed genes in Th2, Th17 and Treg cells include those implicated in relevant biology. For example, Th2 cells are enriched for *GATA3, GATA3-AS1* and *SEMA5A* (Zhang et al., 2013a), *IL17RB* (Wang et al., 2007), *AKAP12* (Lund et al., 2007), *GATA3*, *HPGDS* (Mitson-Salazar et al., 2016) and *PTGDR2* (aka CRTH2). The top ten Th17 reliably expressed genes included *ADAM12* (Zhou et al., 2013), *COL5A3* (Castro et al., 2017), *RORC, CCR6, IL1R1* (Hu et al., 2011), *ABCB1* (Ramesh et al., 2014), *PLXND1* (Guo et al., 2016), and *MAP3K4* (Cleret-Buhot et al., 2015). Finally, the top ten Treg reliably expressed genes known to be important for Treg biology included *CEACAM4* (Hua et al., 2015), *HMCN1* (Sadlon et al., 2010), *DNAH8* (Regateiro et al., 2012), *SEMA3G*, *FOXP3, CSF2RB* (Bhairavabhotla et al., 2016), *IKZF2* (Thornton et al., 2010), *LINC01281* (Ranzani et al., 2015), and *CTLA4* (Takahashi et al., 2000).

Importantly, beyond confirming genes known to be involved in T cell identity, our analysis also revealed genes that may represent novel markers for specific T cell subsets or highlight new biology. For example, *ADAM12, COL5A3, NTN4*, and *TSPAN15* are as at least as consistently expressed in a Th17-specific manner than the commonly used genes *RORC, CCR6*, or *IL17A/IL17F* (Figure 4). *ADAM12* and *TSPAN15* encode for cell-surface-exposed proteins, so could be used for isolating or quantifying Th17 cell populations. Similarly, *CEACAM4, CSF2RB, IKZF2,* and *LINC01281* are as good or better markers at the mRNA level for Treg cells than the commonly used genes *FOXP3, IL2RA, IL10*, or *TGFB*. Again, *CEACAM4* and *CSF2RB* encode for cell-surface-exposed proteins, so may have utility for isolating or quantifying Treg cell populations.

Finally, a major question we sought to answer in this study is whether cytokines, or distinct T cell differentiation programs, also impact alternative splicing. Surprisingly, we find little evidence for widespread coordinated changes in splicing that correlate strongly with T cell subset identity. This could reflect the fact that splicing represents a fine-tuning of T subset function rather than a major determinant. Alternatively, splicing may be more sensitive to variations in the methods used to isolate subpopulation of T cells or heterogeneity inherent in these populations. Regardless, we did identify a few genes for which splicing might contribute to differential function of Treg cells, such as AHRGEF2 and ZNF451. Similar to the unappreciated subset-baised gene expression programs mentioned above, these splicing events represent potentially new biology that we anticipate will motivate further study.

## Acknowledgements

K.W.L. is supported by R35 GM118048, Y.B. is supported by R01 GM128096 and R01AG046544, C.M.R. was supported in part by T32 GM008216, D.B. was supported by a supplement R35 GM118048-S1.

## Author Contributions

C.M.R., Y.B. and K.W.L. designed the study and analysis approach, C.M.R. identified datasets and carried out all the analysis, D.B. isolated cell subsets and generated RNA-Seq libraries, P.J. developed several analysis algorithms used in the study. C.M.R., Y.B. and K.W.L. wrote the manuscript.

## STAR Methods

### Lead Contact and Materials Availability

Further information and requests for resources and reagents should be directed to and will be fulfilled by the Lead Contact, Kristen W Lynch. This study did not generate any new unique reagents.

## EXPERIMENTAL MODEL AND SUBJECT DETAILS

For the in-house Bl dataset, CD4+ human peripheral blood mononuclear cells were obtained via apheresis from de-identified healthy blood donors after informed consent by the University of Pennsylvania Human Immunology Core. Samples were collected from three donors, ND307 (age: 46, sex: Male), ND523 (age: 26, sex: Female), and ND535 (age: 32, sex: Male).

## METHOD DETAILS

### Primary T cell isolation and *in vitro* culturing of Naïve CD4+ T cells into Th0 cells

From the CD4+ T cells apheresis, naïve CD4+ T cells were negatively selected for with MACS Miltenyi CD45RO microbeads (130-046-001). 6-well plates were coated for 3 hours with 2.5 ug anti-CD3 (555336) at 37 degrees Celsius then washed with PBS. 10e6 Naïve CD4+ T cells were then cultured in complete RPMI in the anti-CD3-coated 6 well plates with 2.5 ug soluble anti-CD28 (348040) and 10 IU of IL-2. Cells were harvested after 48 hours.

### RNA-Sequencing of primary T cells

RNA was isolated with RNA Bee (see here for further details: http://www.kwlynchlab.org/s/Isolating-RNA-with-RNABee.pdf) from bulk naïve CD4+ T cells (cultured for 0 hours) and Th0 cells (cultured for 48 hours with anti-CD3 and anti-CD28). The RNA integrity number (RIN) was measured with a bioanalyzer, and all samples had a RIN>8.0. RNA-Sequencing libraries were generated by and sequenced by GeneWiz. The libraries were poly-A selected (non-stranded) and paired-end sequenced at a 150bp read length.

### Selection of datasets and RNA-Seq data processing

#### Database search

The following search terms were used to find appropriate datasets on EMBL-EBI-ArrayExpress (https://www.ebi.ac.uk/arrayexpress/) and NCBI-GEO (https://www.ncbi.nlm.nih.gov/geo/): “T Helper”, “Naïve”, “CD4”, “Th0”, “Th1”, “Th2”,

“Th17”, and “T Regulatory”. Datasets that had at least two of the following types of samples were retained for further analysis: Naïve, Th0, or Th1/Th2/Th17/Treg.

#### RNA-seq data processing steps

SRA files for the publicly available datasets were downloaded from the NCBI Sequence Read Archive. SRA files were converted to fastqs with fastq-dump (sratoolkit.2.9.2) using the following commands: --split-3 –gzip. Sequencing adaptors and low quality base calls were trimmed from reads using trim_galore (0.5.0) using the following commands: --stringency 5 --length 35-q 20. For gene expression analyses, transcript level counts were obtained using Salmon (0.11.3) in mapping-based mode with default settings. The Salmon transcriptome indices were prepared with the GRCh38 genome and Ensembl GRCh38.94 transcript database. Transcript level counts were collapsed into gene level counts with tximport (bioconductor-tximport 1.12.0). For alternative splicing analyses, fastq reads were aligned with STAR (2.5.2a) to the GRCh38 genome supplemented with the Ensembl GRCh38.94 transcript database using the following commands: --outSAMattributes All --alignSJoverhangMin 8 --readFilesCommand zcat -- outSAMunmapped Within. The aligned bam files were then quantified for alternative splicing analyses with MAJIQ (v2.1-59f0404).

#### Identifying and confirming which samples derived from the same human donors

All but one dataset (Ra) used a paired-donor experimental design, meaning two or more samples representing different T cell subsets derived from the same human donor. To determine which samples derived from the same donor, single nucleotide variants were identified for each RNA-Seq sample, and then the proportion of shared genomic variation between samples was calculated. To identify and call the single nucleotide variants from the genome-aligned bam files, bcftools (1.9) was used. The command used was bcftools mpileup -Ou -f <genome.fasta> <bam file> | bcftools call -mv -Ob –output <bcf file>. The bcf files were indexed, and then merged (--output-type z) with bcftools into a vcf file. The merged vcf file was then filtered with bcftools view --min-af 0.25 --output-type z, then the vcf was normalized and converted back to a bcf with bcftools norm -m-any | bcftools norm -Ob --check-ref w -f <genome.fastq>. The resulting bcf was indexed with bcftools, and then processed with plink (v1.90b6.7) using the following commands: --bcf <bcf file> -- const-fid 0 --allow-extra-chr 0 --recode --out <working directory>. To quantify the proportion of shared genomic variation between samples (identify by descent or IBD), plink was run with the following command: --file <working directory> --genome –out <working directory>. Groups of samples that shared greater IBD were determined to derive from the same donor. Reassuringly, this analysis confirmed which samples derived from the same donor in datasets that provided donor information, so we feel confident in in our determination of sample donor pairs for datasets from which donor information was not provided.

#### Quality control checks

The trimmed fastqs were analyzed by FastQC (0.11.2). FastqQC results were used to confirm each sample did not have any non-human overrepresented sequences. One publicly available dataset relevant to this study (but not used for further analyses) failed this quality control check because some samples had over-represented bacterial genome sequences possibly indicating a bacterial contamination during cell culture. To confirm samples from the same donor were convincingly differentiated or sorted into different T cell subtypes, PCA analysis was carried out on the gene-level counts using DESeq2::plotPCA (bioconductor-deseq2 1.22.1). One publicly available dataset relevant to this study (but not used for further analyses) failed this quality control check because the T cell subtype explained less than 10% of the variance in gene expression across samples in the dataset (at least 57% of the variance in gene expression was attributed to the identify of the donor, suggesting inefficient cell sorting). The final quality control check was to confirm a sufficient percentage of reads in the fastqs aligned to the human genome uniquely and unambiguously at a rate of at least 60% according to the STAR logs.

## QUANTIFICATION AND STATISTICAL ANALYSIS

### Differential Expression Analyses

DESeq2 was used to quantify differential expression between T cell subtypes for each dataset. For each test for differential expression between two T cell subtypes, samples were only ever compared from the same dataset. For example, Bl Naïve vs Bl Th0. Before testing for differentially expressed genes, genes were filtered to only retain genes with total counts greater than 10 in at least two samples in at least one subtype. Mitochondrial and ribosomal genes were also filtered out. For all but the Ra and Mi datasets, the donor-aware model matrix supplied to DESeq2 controlled for gene expression variation due to donor by using ‘design = ∼Donor + Subtype’. The Ra and Mi dataset design matrices were simply ‘design = ∼Subtype’. To control for noisy estimates of log2 fold-change in lowly expressed genes and genes with a high coefficient of variation, a log fold-shrinkage was applied to the DESeq2 differential expression results: ‘DESeq2::fcShrink(<deseq_obj>, <coefficient>, type=apelgm, lfcThreshold=1.5, svale=True’).

### Identifying consistently differentially expressed genes between T cell subtypes

To identify core sets of genes highly expressed in each T cell subtypes, we performed differential expression analyses between all pairs of T cell subtypes, controlling for the human donor source of the sample (i.e. Th0 vs Th1, Th2 vs Th1, Th17 vs Th1, TReg vs Th1). Differential expression analyses were done with DESeq2, and the resulting log2 fold changes (log2FC) and the s-values (the probability that the sign of the log2FC is wrong) were used to identify core genes. For example, core Th1 genes were defined as genes more highly expressed (log2FC > 1) in Th1 samples than Th0, Th2, Th17, **or** TReg samples; Th1 core genes needed to be *consistently* higher in Th1 than Th0, Th2, Th17, or TReg in the datasets that included both Th1 and Th0 samples (genes higher in two out two datasets), Th1 and Th2 samples (genes higher in at least four out of five datasets), Th1 and Th17 samples (genes higher in three out of three datasets), or Th1 and TReg samples (genes higher in two out of two datasets). Th1 core gene were further filtered out if none of the differential expression comparisons showed the gene ever having an s-value < 0.001. Core genes were then identified for Th2, Th17, and TReg populations. The resulting core genes are represented in Supplemental Table S3, whereby each core gene can be represented by multiple rows: each row summarizes the differential expression results for the gene from a given T cell comparison (i.e. Th0 vs Th1). In many cases, core genes were identified for multiple T cell subtypes. For example, CXCR3 is a core gene for Th1 (mean log2FC 2.95 over Th2 in five datasets, log2FC 4.2 over Th17 in three datasets) and CXCR3 is also a core gene for TRegs (log2FC 2.36 over Th2 in two datasets).

To better visualize these T cell subtype core genes, the core genes table was filtered, per gene, to select which T cell comparison (i.e. Th0 vs Th1) had the greatest number of datasets showing higher expression with s-value < 0.001 (ties decided by the greatest mean log2FC across the datasets). After filtering, each core gene was represented by a single T cell subtype comparison, and the genes were sorted by which T cell subtype they were more highly expressed in. Figure 4 shows these sorted genes’ asinh TPM levels.

### Splicing Analyses

To look for genes exhibiting consistent differences in splicing between T cell subtypes, the MAJIQ algorithm was used to quantify alternative splicing from the genome-aligned bam files. With the exception of the Ra and Mi datasets, differential splicing was quantified between all pairs of samples from the same donor (e.g. Ka Th1 donor 1 vs Ka Th2 donor 2). For Ra and Mi, samples were compared in bulk versus each other (e.g. all Th1 vs all Th2 in Ra). MAJIQ quantifies the difference in percent splice included for every splice junction (e.g. GeneA junctionX shows a 20% difference in inclusion between Ka Th1 donor 1 vs Ka Th2 donor 2 or 20% difference in inclusion between Ra Th1 samples and Ra Th2 samples). MAJIQ also quantifies the probability that a difference in splice inclusion is above some threshold, and we used a threshold of 20%. Significant differences in splicing were those identified as having a 95% probability of being greater than a 20% difference in splicing.

### Identifying consistently differentially spliced genes between T cell subtypes

To identify consistently differentially spliced splice junctions, for each junction and for each T cell subtype comparison, we first identified the splice junctions that showed a significance difference in splicing in at least one donor from at least one dataset. Next, we filtered out junctions for which any other donors disagreed on the direction of the change in splicing. We next filtered out junctions for which two datasets disagreed on the direction of the change in splicing. Next, we filtered out splicing changes where the number of datasets that agreed the splicing change was at least 10% in the same direction was fewer than two. 43 genes passed these filters, and these genes are listed in Supplemental Table S4.

## DATA AND CODE AVAILABILITY

The new RNA-Seq from naïve and Th0 cells generated for this study is available in GEO (GSE135118). Accessions for the other datasets used in this study include: Hn (E-MTAB-6300), Mi (E-MTAB-5739), Ra (E-MTAB-2319), Tu (GSE52260), Ka (GSE71645), Ab (GSE107981), Re (GSE110097), He (GSE62484), Mo (GSE107011), and Lo (GSE78276). All scripts used to analyze data for this study are made publicly available here: https://bitbucket.org/cradens/t_cell_meta_anlaysis/.

## Supplemental Information

**Supplemental Figure S1:**
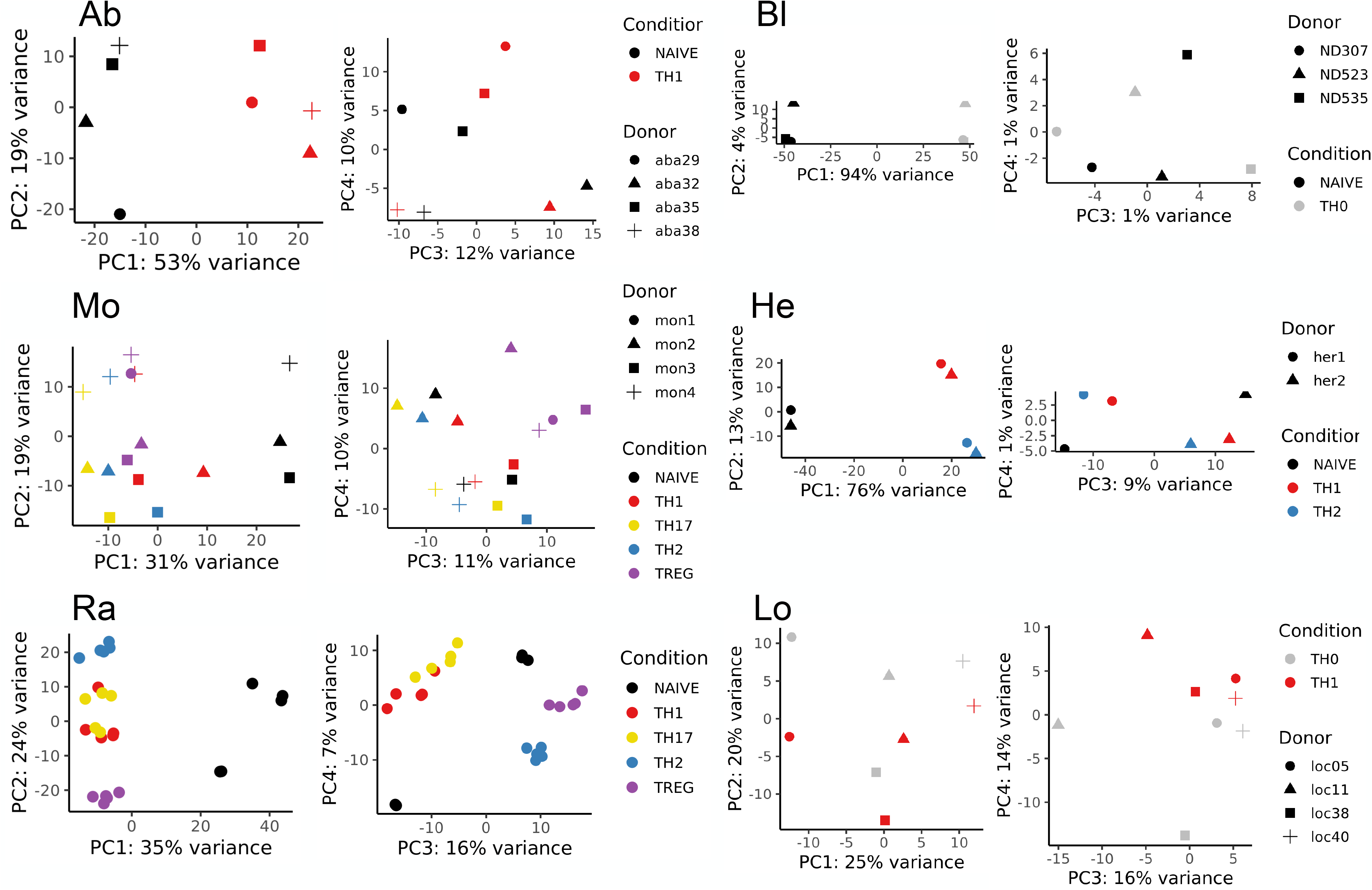
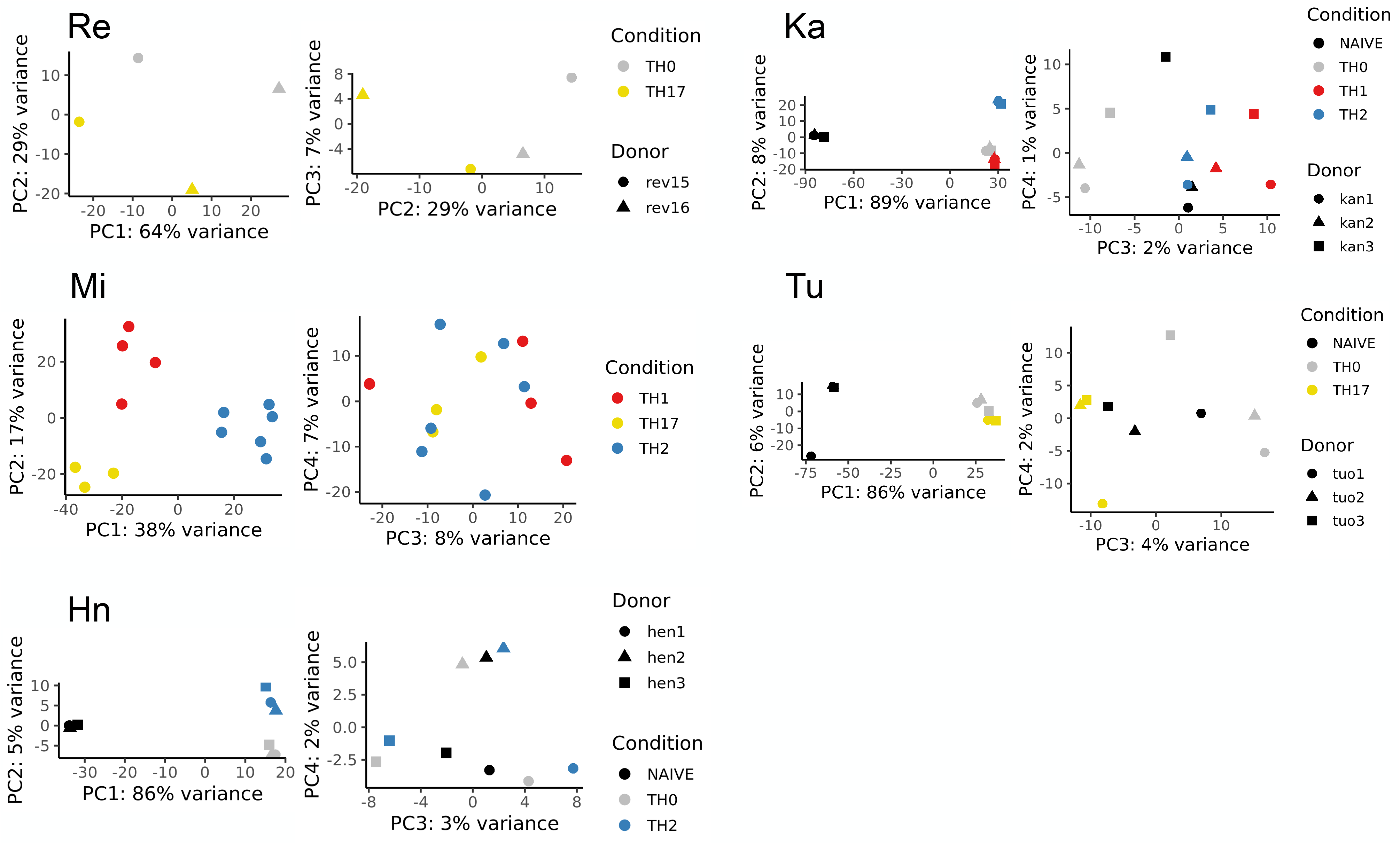
Principle Component Analysis demonstrates that cell types cluster together with each dataset analyzed (related to Figure 2). Principle Component Analysis (PCA) was run on the gene expression values, derived as described in Figure 1c, from RNA-Seq data obtained from 11 distinct studies. Individual samples within each dataset are plotted according to their values of the top two components (PC1, PC2) and colored by T cell subtype. Plots for each dataset are in the same order as in Figure 1A: (a) Ab (ref), (b) Mo (ref), (c) Ra (ref), (d) Bl (this study), (e) He (ref), (f) Lo (ref), (g) Re (ref), (h) Si (ref), (i) Hn (ref), (j) Ka (ref), (k) Tu (ref).

**Supplemental Figure S2:**
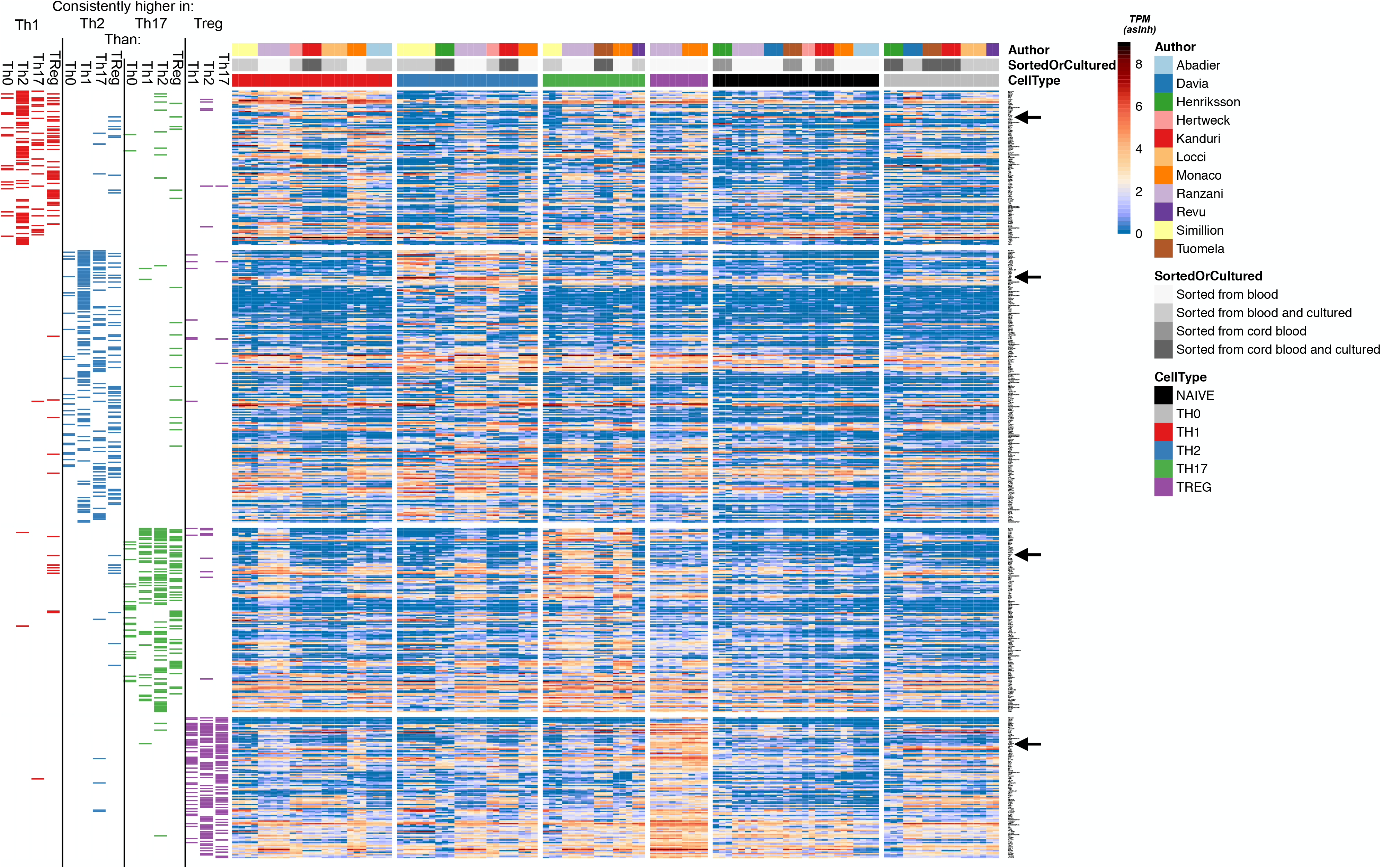
Expression of T subtype-associated cytokine genes across samples analyzed in this study (related to Figure 4). Reliably expressed T Helper genes, ranked by number of supporting datasets and log2FC differences as in Figure 4. Leftmost segment indicates whether a given core T Helper gene was reliably more highly expressed in a given subset over others. All differential expression analyses were performed between samples from the same dataset. The datasets with Treg samples lacked Th0 samples, so there is no “enriched in Treg vs Th0” column. Some T Helper subsets share reliably expressed genes, as indicated by black boxes in multiple columns across a single row. The heatmap shows inverse hyperbolic sine (asinh)-transformed TPMs. Each column is a sample, and samples are grouped by cell type and author. Gene names are on the right. Arrow denotes cut-off of top 20 genes shown in Figure 4.

**Supplemental Figure S3:**
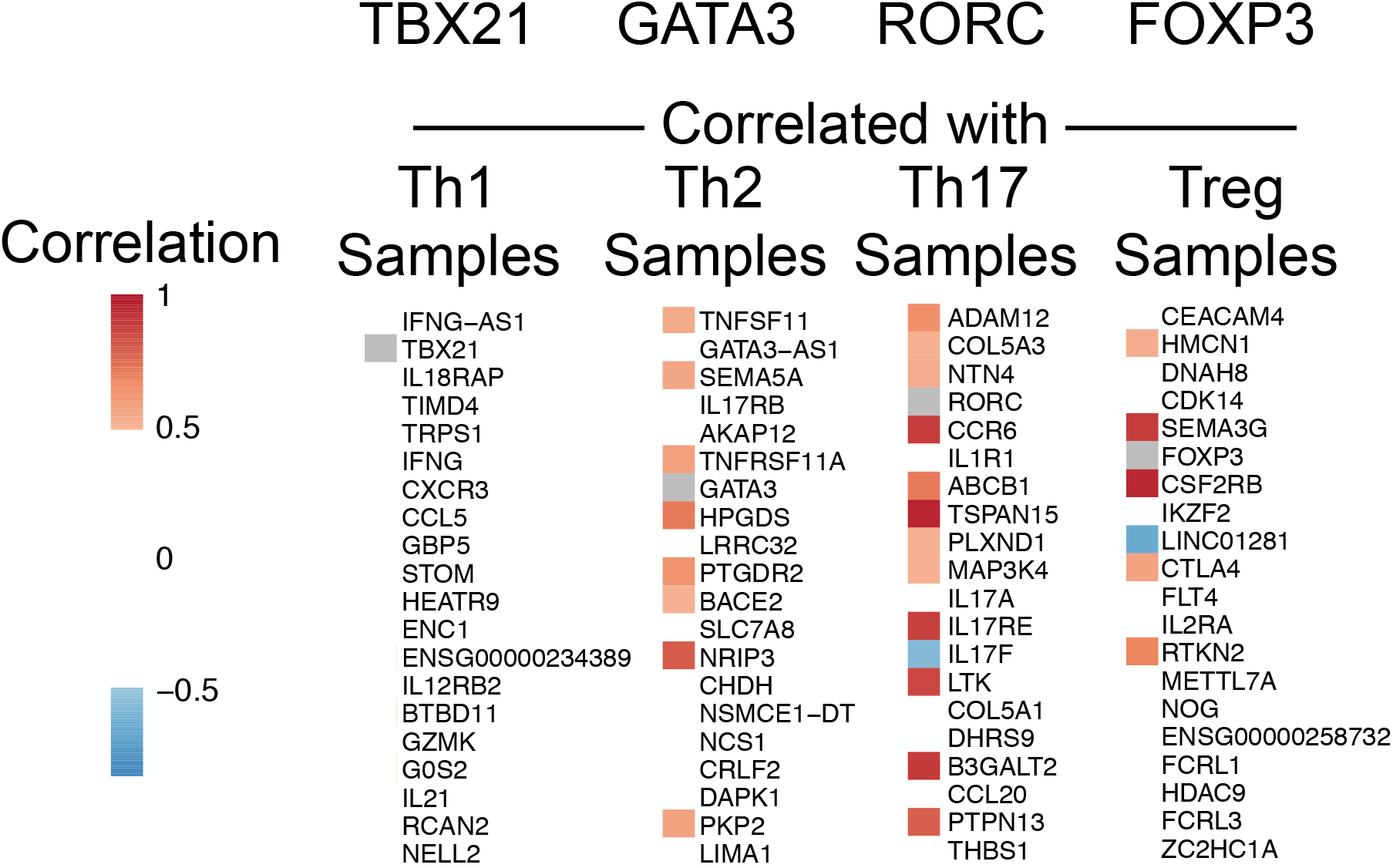
Correlation of expression of master regulator genes with other genes consistently enriched in the subtype (related to Figure 4). Gene expression was correlated in a pairwise fashion between master regulator (top of column) and individual subtype-biased genes. Colors indicate the correlation value as indicated in key.

**Supplemental Table S1: (Related to Figure 1) RNA-Sequencing technical specifications.**

**Supplemental Table S2: (Related to Figure 1) Purification methods for cell populations used in this study**

**Supplemental Table S3: (related to Figure 4) Genes consistently expressed in each T cell subset**

**Supplemental Table S4: (related to Figure 5) Genes exhibiting consistent alternative splicing patterns in each T cell subset**

---

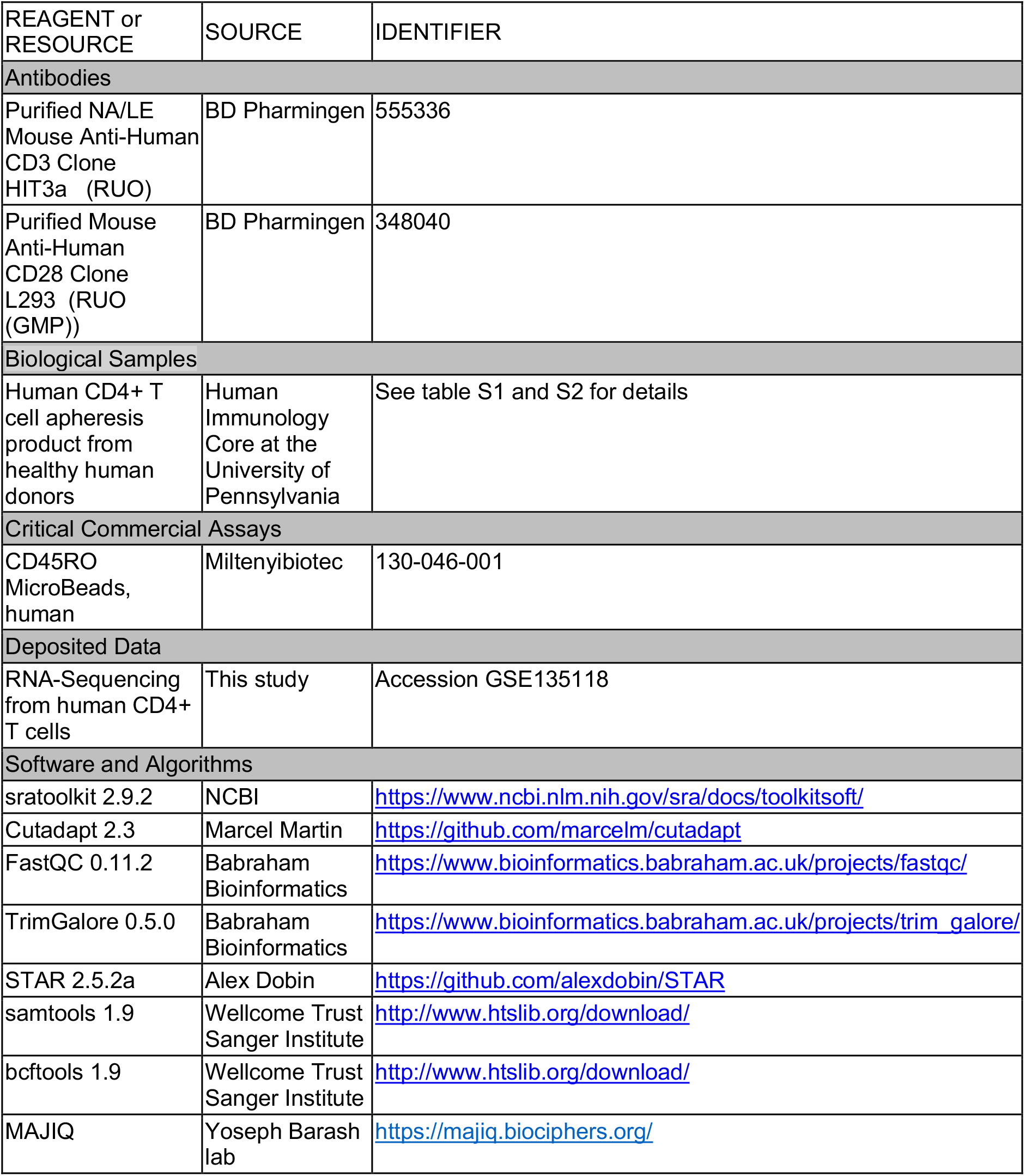
KEY RESOURCES TABLE.

